# PsychRNN: An Accessible and Flexible Python Package for Training Recurrent Neural Network Models on Cognitive Tasks

**DOI:** 10.1101/2020.09.30.321752

**Authors:** Daniel B. Ehrlich, Jasmine T. Stone, David Brandfonbrener, Alexander Atanasov, John D. Murray

**Affiliations:** Interdepartmental Neuroscience Program, Yale University, New Haven, CT, USA; Department of Computer Science, Yale University, New Haven, CT, USA; Department of Computer Science, New York University, New York, NY, USA; Department of Physics, Yale University, New Haven, CT, USA; Department of Physics, Harvard University, Cambridge, MA, USA; Department of Psychiatry, Yale School of Medicine, New Haven, CT, USA

## Abstract

Task-trained artificial recurrent neural networks (RNNs) provide a computational modeling framework of increasing interest and application in computational, systems, and cognitive neuroscience. RNNs can be trained, using deep learning methods, to perform cognitive tasks used in animal and human experiments, and can be studied to investigate potential neural representations and circuit mechanisms underlying cognitive computations and behavior. Widespread application of these approaches within neuroscience has been limited by technical barriers in use of deep learning software packages to train network models. Here we introduce PsychRNN, an accessible, flexible, and extensible Python package for training RNNs on cognitive tasks. Our package is designed for accessibility, for researchers to define tasks and train RNN models using only Python and NumPy without requiring knowledge of deep learning software. The training backend is based on TensorFlow and is readily extensible for researchers with TensorFlow knowledge to develop projects with additional customization. PsychRNN implements a number of specialized features to support applications in systems and cognitive neuroscience. Users can impose neurobiologically relevant constraints on synaptic connectivity patterns. Furthermore, specification of cognitive tasks has a modular structure, which facilitates parametric variation of task demands to examine their impact on model solutions. PsychRNN also enables task shaping during training, or curriculum learning, in which tasks are adjusted in closed-loop based on performance. Shaping is ubiquitous in training of animals in cognitive tasks, and PsychRNN allows investigation of how shaping trajectories impact learning and model solutions. Overall, the PsychRNN framework facilitates application of trained RNNs in neuroscience research.

**Visual Abstract:** Example workflow for using PsychRNN. First, the task of interest is defined, and a recurrent neural network model is trained to perform the task, optionally with neurobiologically informed constraints on the network. After the network is trained, the researchers can investigate network properties including the synaptic connectivity patterns and the dynamics of neural population activity during task execution, and other studies, e.g. those on perturbations, can be explored. The dotted line shows the possible repetition of this cycle with one network, which allows investigation of training effects of task shaping, or curriculum learning, for closed-loop training of the network on a progression of tasks.

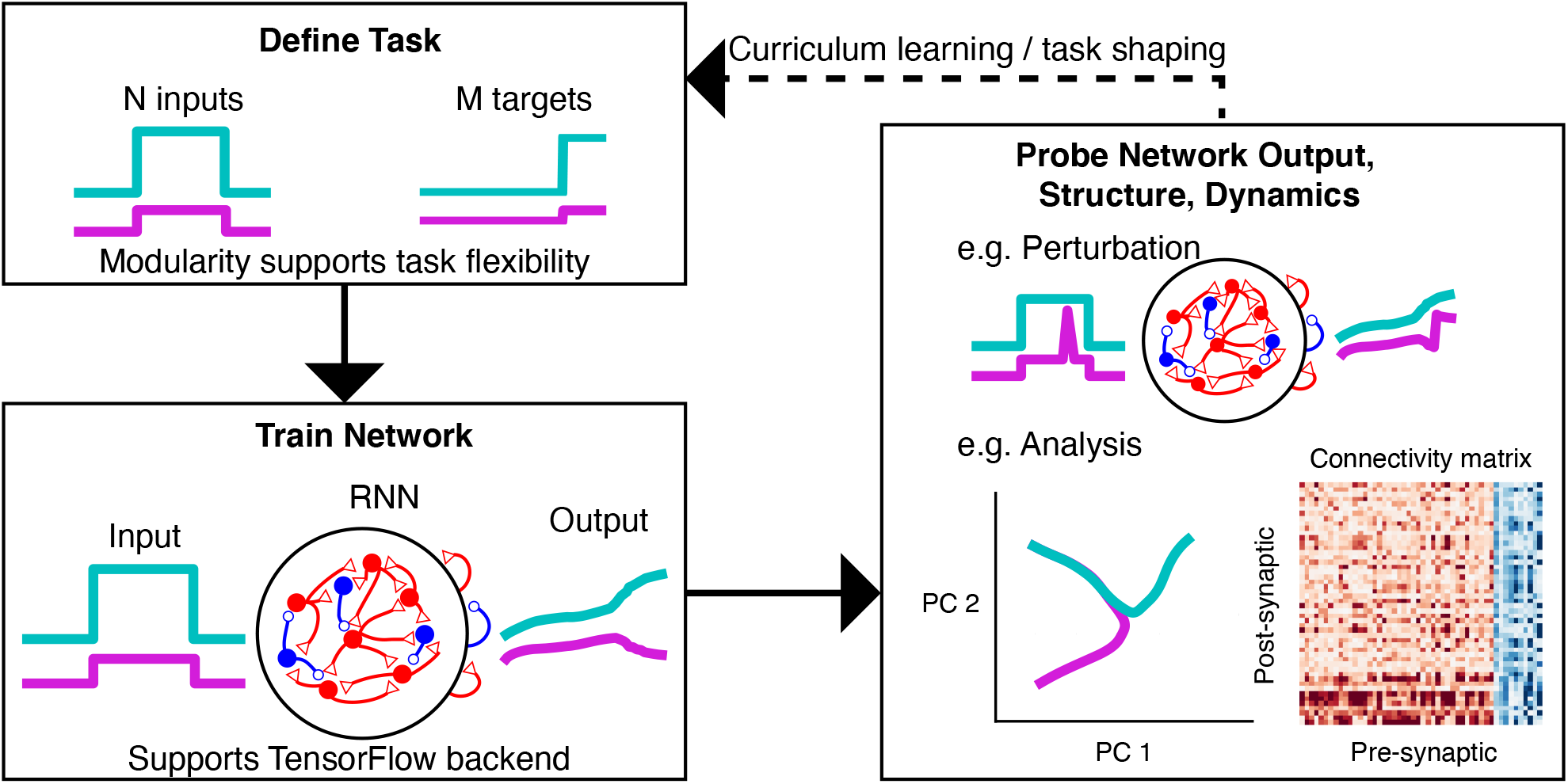

**Significance Statement:** Artificial recurrent neural network (RNN) modeling is of increasing interest within computational, systems, and cognitive neuroscience, yet its proliferation as a computational tool within the field has been limited due to technical barriers in use of specialized deep-learning software. PsychRNN provides an accessible, flexible, and powerful framework for training RNN models on cognitive tasks. Users can define tasks and train models using the Python-based interface which enables RNN modeling studies without requiring user knowledge of deep learning software. PsychRNN’s modular structure facilitates task specification and incorporation of neurobiological constraints, and supports extensibility for users with deep learning expertise. PsychRNN’s framework for RNN modeling will increase accessibility and reproducibility of this approach across neuroscience subfields.

## Introduction

Studying artificial neural networks (ANNs) as models of brain function is an approach of increasing interest in computational, systems, and cognitive neuroscience (Kriegeskorte, 2015; Yamins and DiCarlo, 2016; Richards et al., 2019). ANNs comprise many simple units, called neurons, whose synaptic connectivity patterns are iteratively updated via deep learning methods to optimize an objective. For application in neuroscience and psychology, ANNs can be trained to perform a cognitive task of interest, and the trained networks can then be analyzed and compared to experimental data in a number of ways, including their behavioral responses, neural activity patterns, and synaptic connectivity. Recurrent neural networks (RNNs) form a class of ANN models which are especially well-suited to perform cognitive tasks which unfold across time, common in psychology and neuroscience, such as decision-making or working-memory tasks (Sussillo, 2014; Song et al., 2016; Barak, 2017; Yang and Wang, 2020). In RNNs, highly recurrent synaptic connectivity is optimized to generate target outputs through the network population dynamics. RNNs have been applied to model the dynamics of neuronal populations in cortex during cognitive, perceptual, and motor tasks and are able to capture associated neural response dynamics (e.g., Mante et al., 2013; Sussillo et al., 2015; Carnevale et al., 2015; Rajan et al., 2016; Remington et al., 2018; Masse et al., 2019).

Despite growing impact of RNN modeling in neuroscience, wider adoption by the field is currently hindered by the requisite knowledge of specialized deep learning platforms, such as TensorFlow or PyTorch, to train RNN models. This creates accessibility barriers for researchers to apply RNN modeling to their neuroscientific questions of interest. It can be especially challenging in these platforms to implement neurobiologically motivated constraints, such as structured synaptic connectivity or Dale’s principle defining excitatory and inhibitory neurons. There is also need for modular frameworks to define the cognitive tasks on which RNNs are trained, which would facilitate investigation of how task demands shape network solutions. To better model experimental paradigms for training animals on cognitive tasks, an RNN framework should enable investigation of task shaping, in which training procedures are progressive adapted to the subject’s performance during training.

To address these challenges, we developed the software package PsychRNN as an accessible, flexible, and extensible computational framework for training RNNs on cognitive tasks. Users define tasks and train RNN models using only Python and NumPy, without requiring knowledge of deep learning software. The training backend is based on TensorFlow and is extensible for projects with additional customization. PsychRNN implements a number of specialized features to support applications in systems and cognitive neuroscience, including neurobiologically relevant constraints on synaptic connectivity patterns. Specification of cognitive tasks has a modular structure, which aids parametric variation of task demands to examine their impact on model solutions and promotes code reuse and reproducibility. Modularity also enables implementation of curriculum learning, or task shaping, in which tasks are adjusted in closed-loop based on performance. Our overall goal for PsychRNN is to facilitate application of RNN modeling in neuroscience research.

## Materials and Methods

### Package structure

To serve our objectives of accessibility, extensibility, and reproducibility, we divided the PsychRNN package into two main components: the `Task` object and the Backend (**Fig. 1**). We anticipate that all PsychRNN users will want to be able to define novel tasks specific to their research domains and questions. The `Task` object is therefore fully accessible to users without any TensorFlow or deep learning background. Users familiar with Python and NumPy are able to fully customize novel tasks, and they can customize network structure (e.g., number of units, form of nonlinearity, connectivity) through preset options built into the Backend.

**Figure 1:**
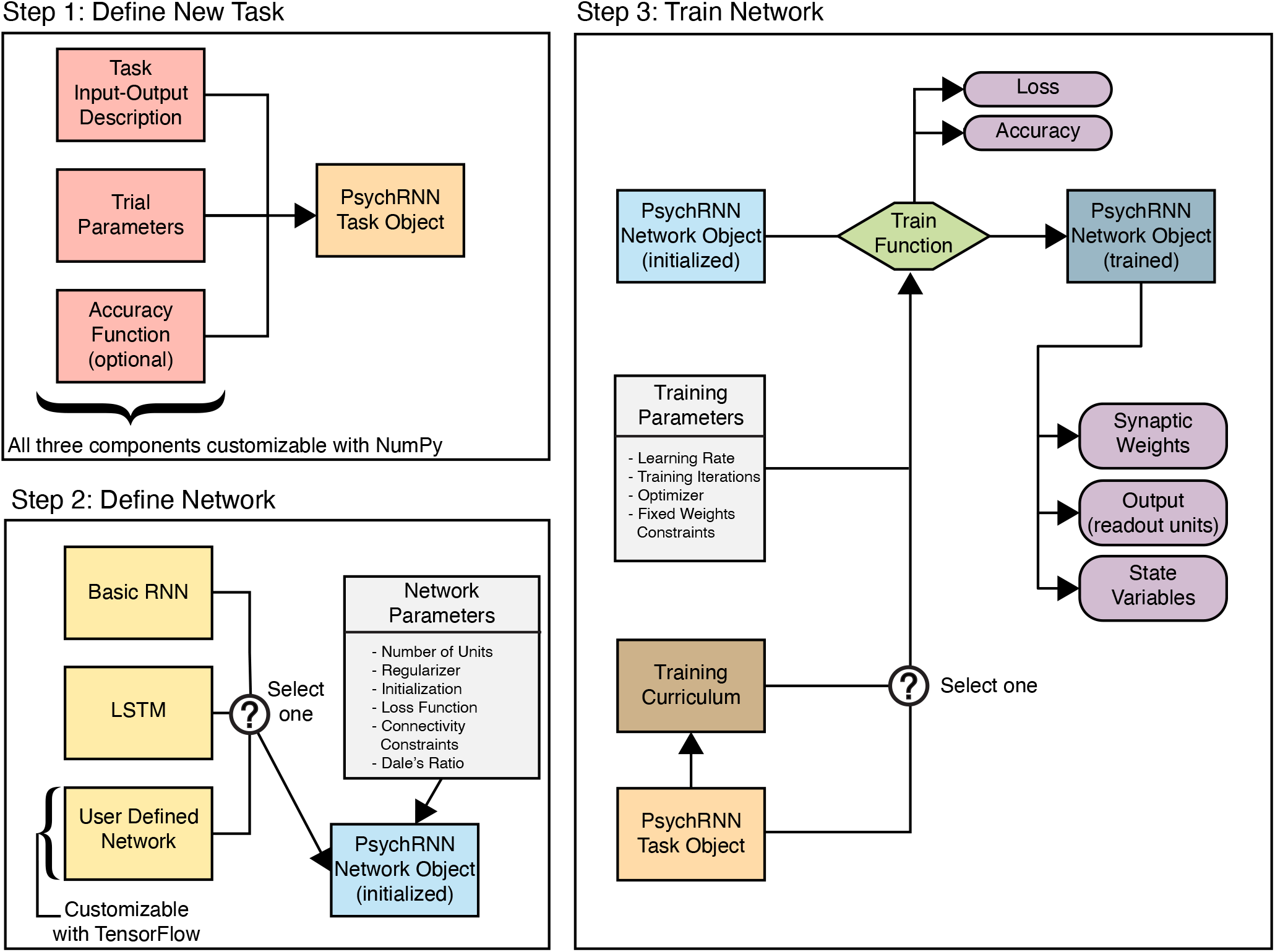
PsychRNN package structure. (**Step 1**) Defining a new task requires two NumPy-based components: `trial-function` describes the task inputs and outputs, and `generate_trial_params` defines parameters for a given trial (**Extended Data, Figure 1-1**). Optionally, one can define an accuracy function describing how to calculate whether performance on a trial was successful. **(Step 2)** The Backend defines the network. First, the model, or network architecture, is selected. A basic RNN and LSTM (Hochreiter and Schmidhuber, 1997) are implemented, and more models or architectures can be defined using TensorFlow. That model is then instantiated with a dictionary of parameters, which includes the number of recurrent units and may also include specifications of loss functions, initializations, regularizations, or constraints. If any parameter is not set, a default is used. **(Step 3)** Training specifications, such as the optimizer or curriculum, can be specified. During network training, measures of performance (loss and accuracy) are recorded at regular intervals. Optimization of the network weights is performed to minimize the loss. After training, the synaptic weight matrix can be saved, and state variables and network output can be generated for any given trial.

For users with greater need for flexibility in network design, the Backend is designed for accessibility, customizability, and extensibility. Backend customization typically requires knowledge of TensorFlow. For those with TensorFlow knowledge, PsychRNN’s modular design enables definition of new models, regularizations, loss functions, and initializations. This modularity facilitates testing hypotheses regarding the impact of specific potential structural constraints on RNN training without having to expend time and resources designing a full RNN codebase.

### Task object

The `Task` object is structured to allow users to define their own new task using Python and NumPy. Specifically, `generate_trial_params` creates trial specific parameters for the task (e.g., stimulus and correct response). `trial_function` specifies the input, target output, and output mask at a given time `t`, given the parameters generated by `generate_trial_params`. PsychRNN comes set with three example tasks that are well researched by cognitive neuroscientists: perceptual discrimination (Roitman and Shadlen, 2002), delayed discrimination (Romo et al., 1999), and delayed match-to-category (Freedman and Assad, 2006). These tasks highlight possible schemas users can apply to specifying their own tasks and provide tasks with which users can test the effect of different structural network features.

Tasks can optionally include `accuracy` functions. Accuracy measures performance in a manner more relevant to experiments than traditional machine learning measures such as loss. On a given trial, accuracy is either one (success) or zero (failure). In contrast, loss on a given trial is a real-numbered value. Accuracy is calculated over multiple trials to obtain a ratio of trials correct to total trials. Accuracy is used as the default metric by the `Curriculum` class.

### Backend

The Backend includes all of the neural network training and specification details (**Fig. 1, Step 2**). The backend, while being accessible and customizable, was designed with pre-set defaults sufficient to get started with PsychRNN. The TensorFlow details are abstracted away by the Backend so that researchers are free to work with or without an understanding of TensorFlow. Additionally, since the Backend is internally modular, different components of the Backend can be swapped in and out interchangeably. In this section, modular components of the Backend are described so that researchers who want to get more in-depth with PsychRNN know what tools are available to them.

### Models

RNNs are a large class of ANNs that process input over time. In the PsychRNN release, we include a basic RNN (which we refer to as an RNN throughout the rest of the paper), and an LSTM model (Hochreiter and Schmidhuber, 1997). The basic RNN model is governed by the following equations:

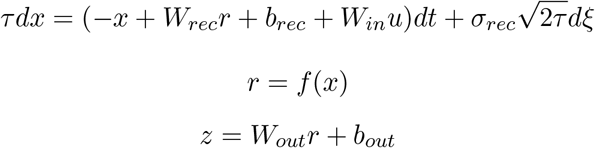

where *u, x* and *z* are the input, recurrent state, and output vectors, respectively. *W_in_*, *W_rec_* and *W_out_* are the input, recurrent, and output synaptic weight matrices. *b_rec_* and *b_out_* are constant biases into the recurrent and output units. *dt* is the simulation time-step and *τ* is the intrinsic timescale of recurrent units. *σ_rec_* is a constant to scale recurrent unit noise, and *dξ* is a gaussian noise process with mean 0 and standard deviation 1. *f*(*x*) is a nonlinear transfer function, which by default in PsychRNN is rectified linear (ReLU). This default can be replaced with any TensorFlow transfer function.

PsychRNN also includes an implementation of LSTMs (Long Short Term Memory networks), a special class of RNNs that enables longer-term memory than is easily attainable with basic RNNs (Hochreiter and Schmidhuber, 1997). LSTMs use a separate “cell state” to store information gated by sigmoidal units. Additional models can be user-defined but require knowledge of TensorFlow.

### Initializations

The synaptic weights that define an ANN are typically initialized randomly. However, with RNNs, large differences in performance, training time and total asymptotic loss, have been observed for different initializations (V. Le et al., 2015). Since initializations can be crucial for training, we have included several initializations currently used in the field (Glorot and Bengio, 2010). By default, recurrent weights are initialized randomly from a gaussian distribution with spectral radius of 1.1 (Sussillo and Abbott, 2009). We also include an initialization called Alpha Identity that initializes the recurrent weights as an identity matrix scaled by a parameter *α* (V. Le et al., 2015). Each of these initializations can substantially improve the training process of RNNs. PsychRNN includes a `Weightinitialization` class that initializes all network weights randomly, all biases as zero, and connectivity masks as all-to-all. New initializations inherit this class and can override any variety of initializations defined in the base class.

### Loss Functions

During training an RNN is optimized to minimize the loss, so the choice of loss function can be crucial for determining exactly what the network learns. By default, the loss function is `mean_squared_error`. Our Backend also includes an option for using `binary_cross_entropy` as the loss function. Other loss functions can be easily defined with some TensorFlow knowledge and added to the `LossFunction` class. Loss functions take in the network output (`predictions`), the target output (`y`) and the output mask, and return a float calculated using the TensorFlow graph.

### Regularizers

Regularizers are penalties added to the loss function that may help prevent the network from overfitting to the training data. We include options for L2-norm and L1-norm regularization for the synaptic weights, which tend to reduce the magnitude of weights and sparsify the resulting weight matrices. In addition we include L2-norm regularization on the post-nonlinearity recurrent unit activity *r*. Other regularizations can be added to the `Regularizer` class through TensorFlow. By default, no regularizations are used.

### Optimizers

PsychRNN is built to take advantage of the many optimizers available in the TensorFlow package. Instead of explicitly defining equations for back propagation through time (BPTT), PsychRNN converts the user supplied Task and RNN into a “graph” model interpretable by TensorFlow. TensorFlow can then automatically generate gradients of the user supplied `LossFunction` with respect to the weights of the network. These gradients can then be used by any TensorFlow optimization algorithm such as stochastic gradient descent, Adam or RMSProp to update the weights and improve task performance (Ruder, 2017).

### Neurobiologically motivated connectivity constraints

PsychRNN is designed for investigation of neurobiologically motivated constraints on the input, recurrent, and output synaptic connectivity patterns. The user can specify which synaptic connections are allowed and which are forbidden (set to zero) through optional user-defined masks at the point of RNN model initialization. This feature enables modeling neural architectures including sparse connectivity and multi-region networks (Rikhye et al., 2018). Optional user-defined masks allow specification of which connections are fixed in their weight values, and which connections are plastic for optimization during training (Rajan et al., 2016). By default, all weights are allowed to be updated by training. PsychRNN also enables implementation of Dale’s principle, such that each recurrent unit’s synaptic weights are all of the same sign (i.e., each neuron’s post-synaptic weights are either all excitatory or all inhibitory) (Song et al., 2016). The optional parameter `dales_ratio` sets the proportion of excitatory units, making the balance inhibitory.

### Curriculum learning

Curriculum learning refers to the presentation of training examples structured into successive discrete blocks sorted by increasing difficulty (Bengio et al., 2009; Krueger and Dayan, 2009). Task modularity in PsychRNN enables an intuitive framework for curriculum learning that does not require TensorFlow knowledge. Curriculum learning is implemented by passing a `Curriculum` object to the RNN model when training is executed. Although very flexible and customizable, the simplest form of the `Curriculum` object can be instantiated solely with the list of tasks that one wants to train on sequentially.

The `Curriculum` class included in PsychRNN is flexible and extensible. By default, accuracy, as defined within a task, is used to measure the performance of the task. When the performance surpasses a user-defined threshold, the network starts training with the next task. The `Curriculum` object thus includes an optional input array, `thresholds`, for specifying the performance thresholds required to advance to each successive task. Apart from accuracy one may wish to advance the curriculum stage using an alternative measure such as loss or number of iterations. We include an optional `metric` function that can be passed into the `Curriculum` class to define a custom measure to govern task stage transitions.

### Simulator

One limitation of specifying RNN networks in the TensorFlow language is that in order to run a network the inputs, outputs, and computation need to take place within the TensorFlow framework, which can impede users’ ability to design and implement experiments on their trained RNN models. To mitigate this, we have included a NumPy-based simulator which takes in RNN and `Task` objects and simulates the network in NumPy. This simulator allows the user to study various neuroscientific topics such as robustness to perturbations.

### Software availability

The PsychRNN open-source software described in the paper is available online for download in a Git-based repository at https://github.com/murraylab/PsychRNN. Detailed documentation containing tutorials and examples is also provided. The code and documentation are available as Extended Data. All data and figures included were produced on a MacBook Pro (Retina, 13-inch, Early 2015) with 8 GB of RAM and 2.7 GHz running macOS Catalina 10.15.5 in an Anaconda environment with Python 3.6.9, NumPy 1.17.2, and TensorFlow 1.14.0.

## Results

To facilitate accessibility, PsychRNN allows users to define tasks and define and train networks using a Python- and NumPy-based interface. PsychRNN provides a machine-learning backend, based on TensorFlow, which converts task and network specifications into the Tensorflow deep learning framework to optimize network weights. This allows users to focus on the neuroscientific questions rather than implementation details of deep learning software packages. As an example, we demonstrate how PsychRNN can specify an RNN model, train it to perform a task of neuroscientific interest—here, a two-alternative forced-choice perceptual discrimination task (Roitman and Shadlen, 2002)—and return behavioral readout from output units and internal activity patterns of recurrent units (**Fig. 2**).

**Figure 2:**
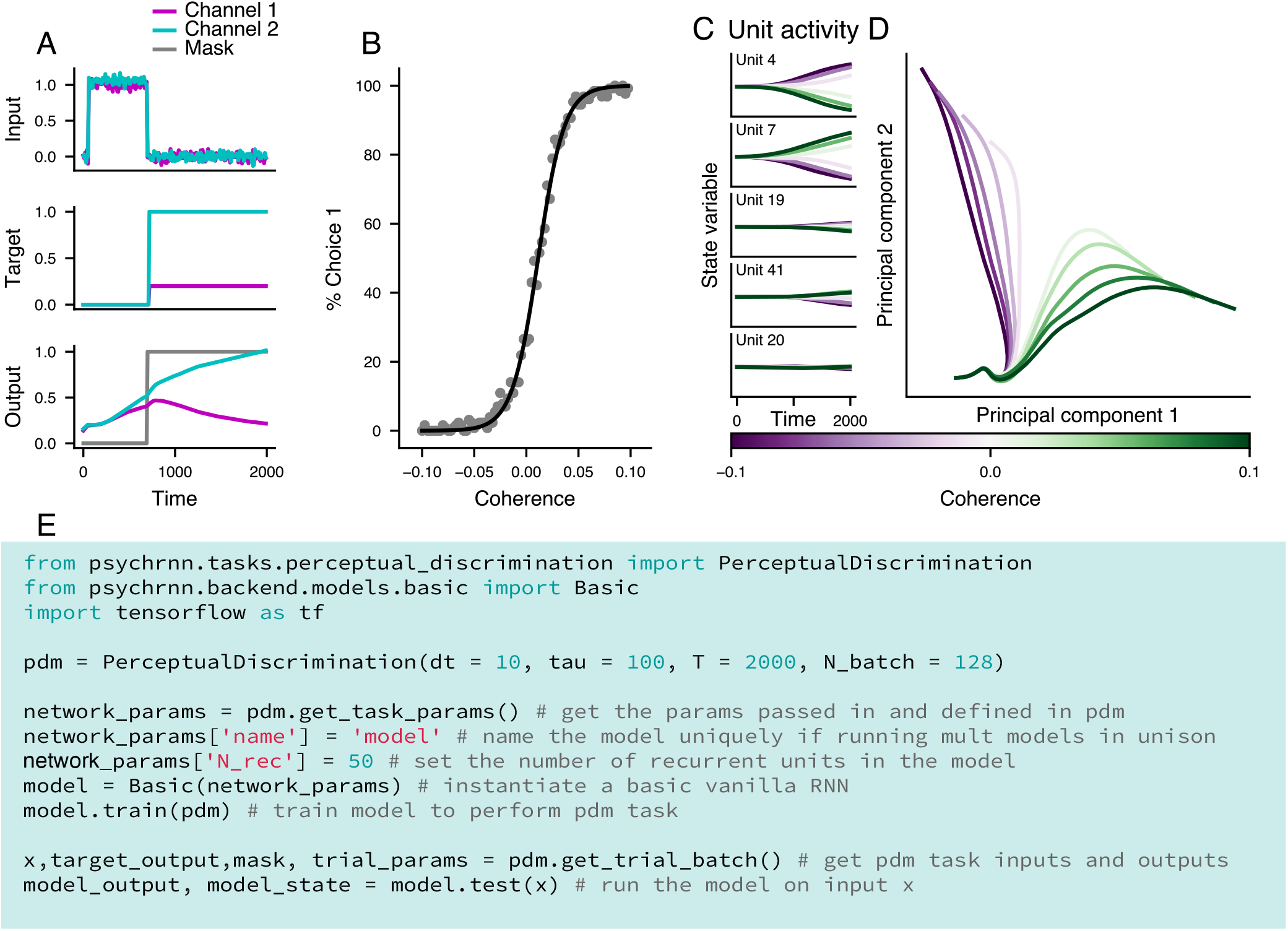
Example Task (Perceptual Discrimination). **(A)** Inputs and target output as specified by the task (top two panels), and the network’s output for the displayed input (bottom panel). Because the output mask is zero during the stimulus period, the network is not directly constrained during that period. **(B)** Percent of decisions the network makes for Choice 1 at varying coherence levels. Negative coherences levels indicate stimulus inputs rewarded Choice 2. A psychometric function is fit to the data (black). This plot validates that the network successfully learned the task. **(C)** State variable activity traces across for a range of stimulus coherences, for multiple example units, averaged over correct trials. The network produces state variable activity across all units. **(D)** Population activity traces in the subspace of the top two principal components (PCs). PCA was applied to the activity matrix formed by concatenating across coherences the trialaveraged correct-trial traces, for each unit. **(E)** Minimal example code for using PsychRNN. All relevant modules are imported (lines 1-4), a PerceptualDiscrimination `Task` object is initialized (lines 6-11), the basic RNN model is initialized, built, and trained (lines 13-15), output and state variables are extracted (lines 17-18).

### Modularity

The PsychRNN Backend is complimented by the `Task` object which enables users to easily and flexibly specify tasks of interest without any prerequisite knowledge of TensorFlow or machine learning. The `Task` object allows flexible input and output structure, with tasks varying in not only the task-specific features but also the number of input and output channels (**Fig. 3**). Furthermore, the object-oriented structure of task definition in PsychRNN facilitates tasks that can be quickly and easily varied along multiple dimensions. For example in an implementation of a delayed discrimination task (Romo et al., 1999), we can vary stimuli and delay durations with a set of two parameters (**Fig. 3B**). Importantly not only can we vary the inputs as they exist, but integration between the Task object and Backend makes it possible to vary the structure of the network from the Task object. In our implementation of a delayed match-to-category task (Freedman and Assad, 2006), we can freely change the number of inputs (input discretization) and the number of outputs (categories) (**Fig. 3D**). This flexibility allows researchers to investigate how the network solution of trained RNNs may depend on task or structural properties (Orhan and Ma, 2019).

**Figure 3:**
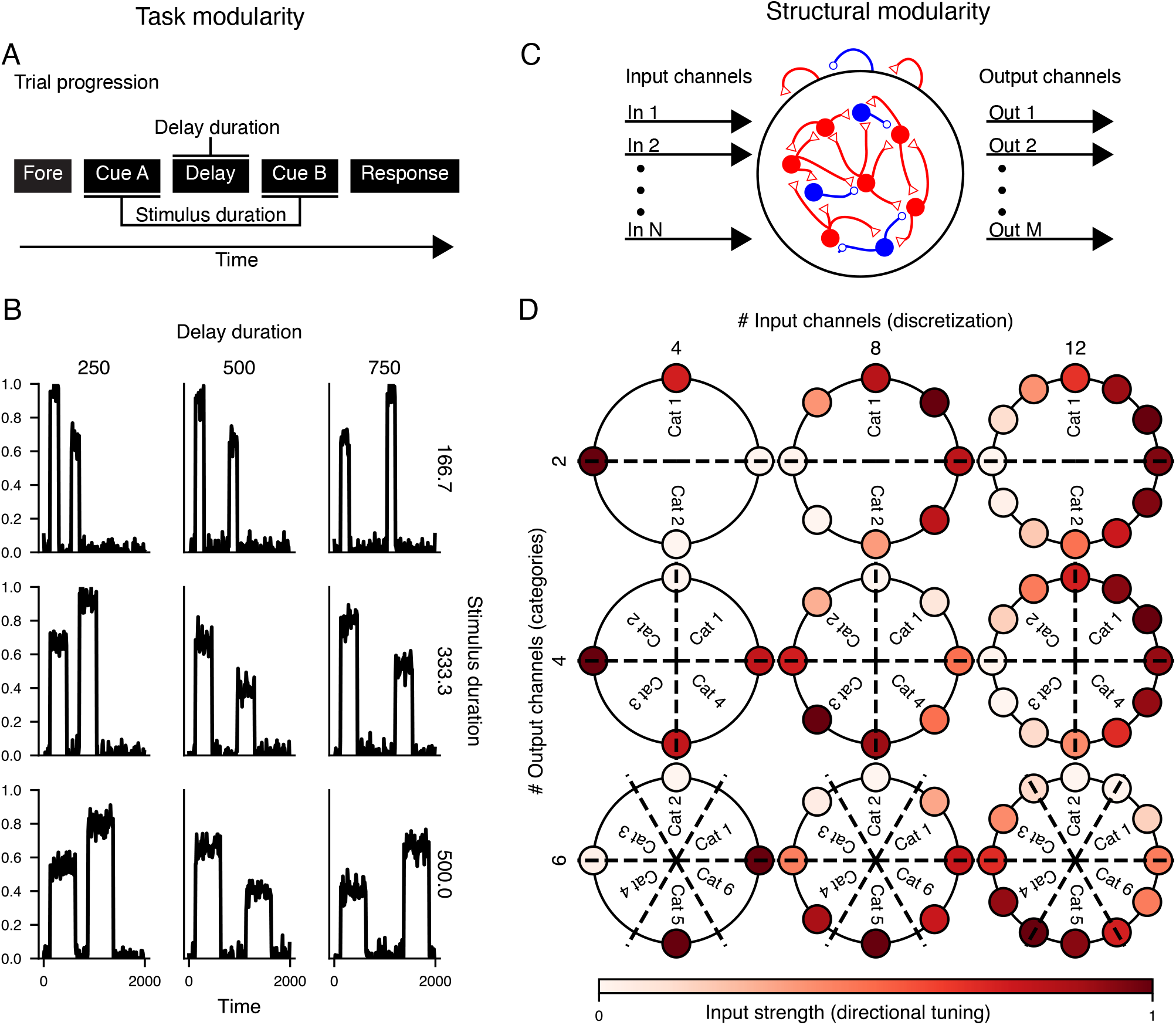
Modularity of task definition. **(A)** Task modularity. This schematic illustrates the trial progression of one trial of a delayed discrimination task. The task is modularly defined such that stimulus and delay duration can be varied easily, simply by changing task parameters. **(B)** One input channel generated by a delayed discrimination task, with varied stimulus and delay durations (**Extended Data, Figure 3-1**). Delay duration is varied across columns, and stimulus duration is varied across rows. **(C)** Structural modularity. Tasks can provide any numbers of channels for input and output on which to train a particular RNN model. Variation in numbers of inputs and outputs is enabled through simple modular task parameters in PsychRNN. **(D)** Example of a match-to-category task. The number of inputs (colored outer circles) is varied across columns, and the number of output categories (Cat) is varied across rows (**Extended Data, Figure 3-2**).

### Neurobiologically motivated connectivity constraints

While there are multiple general-purpose frameworks for training ANNs, neuroscientific modeling often requires neurobiologically motivated constraints and processes which are not common in general-purpose ANN software. PsychRNN includes a variety of easily implemented forms of constraints on synaptic connectivity. The default RNN network has all-to-all connectivity, and allows units to have both excitatory and inhibitory connections. Users can specify which potential synaptic connections are forbidden or allowed, as well as which are fixed and which are plastic for updating during training. Furthermore, PsychRNN can enforce Dale’s principle, so that each unit has either all-excitatory or all-inhibitory synapses onto its targets. **Fig. 4** demonstrates that networks can be trained while subject to various constraints on recurrent connectivity. For example, units can be prevented from making autapses (i.e., self-connections). Networks with block-like connectivity matrices can be used to model multiple brain regions, with denser within-region connectivity and sparser between-region connectivity.

**Figure 4:**
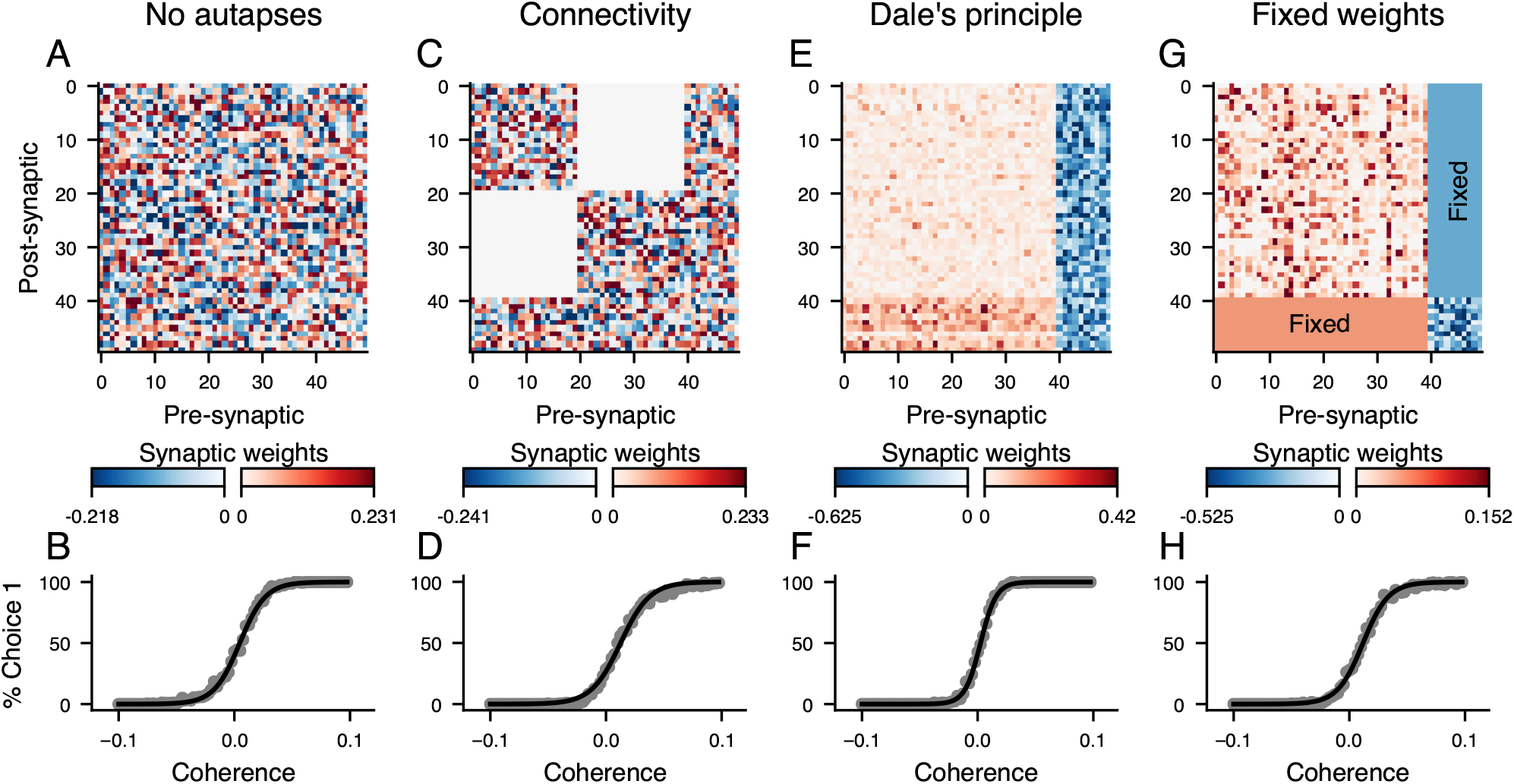
Neurobiologically motivated constraints. This figure illustrates the effects of different connectivity constraints on the recurrent weight matrices and psychometric functions of RNNs trained on the perceptual discrimination task (**Figure 2**). For the recurrent weight matrices (top row), red and blue show excitatory and inhibitory connections, respectively. The coherence plots (bottom row) show that the network successfully trains to perform the task while adhering to the constraints. **(A,B)** This network is constrained to have no autapses, i.e., no self-connections, as illustrated by zeros along the diagonal of the weight matrix. **(C,D)** This network is constrained to have two densely connected populations of units with sparse connection between the populations. This constraints can be used to simulate long-range interactions among different brain regions. **(E,F)** This network is constrained to follow Dale’s principle: each neuron either has entirely excitatory or entirely inhibitory outputs. **(E,F)** This network has Dale’s principle enforced and has subset of weights which are fixed, i.e., they cannot be updated by training. In this example, all connections between excitatory and inhibitory neurons are fixed, and other within excitatory-to-excitatory and inhibitory-to-inhibitory connections are plastic during training.

### Curriculum learning

One important feature included in PsychRNN is a native implementation of curriculum learning. Curriculum learning, also referred to as task shaping in the psychological literature (Krueger and Dayan, 2009), refers to structuring training examples such that the agent learns easier trials or more basic subtasks first (**Fig. 5A,B**). Curriculum learning has been shown to improve artificial neural network training both in training iterations to convergence and in the final loss (Bengio et al., 2009). In neuroscience, researchers adopt a wide variety of different curricula to train animals to perform full experimental tasks. By including curriculum learning, PsychRNN enables researchers to investigate how training curricula may impact resulting behavioral and neural solutions to cognitive tasks, as well as potentially identify new curricula that may accelerate training. Further, curricula can be used more broadly to investigate how learning may be influenced and biased by the sets of tasks an agent has previously encountered. Lastly, curriculum learning can be used to train networks on tasks that may be too complex to be learned without it. As an example, we trained RNNs on a version of the perceptual decision-making task (from **Fig. 2**), and examined the effects of using curriculum learning in the training procedure (**Fig. 5C,D**). Here, curriculum learning involved initially training the model at high stimulus coherences, and introducing progressively lower coherences when the model’s performance reached a threshold level. We found that curriculum learning enabled faster training of models, as commonly observed in experiments (Krueger and Dayan, 2009).

**Figure 5:**
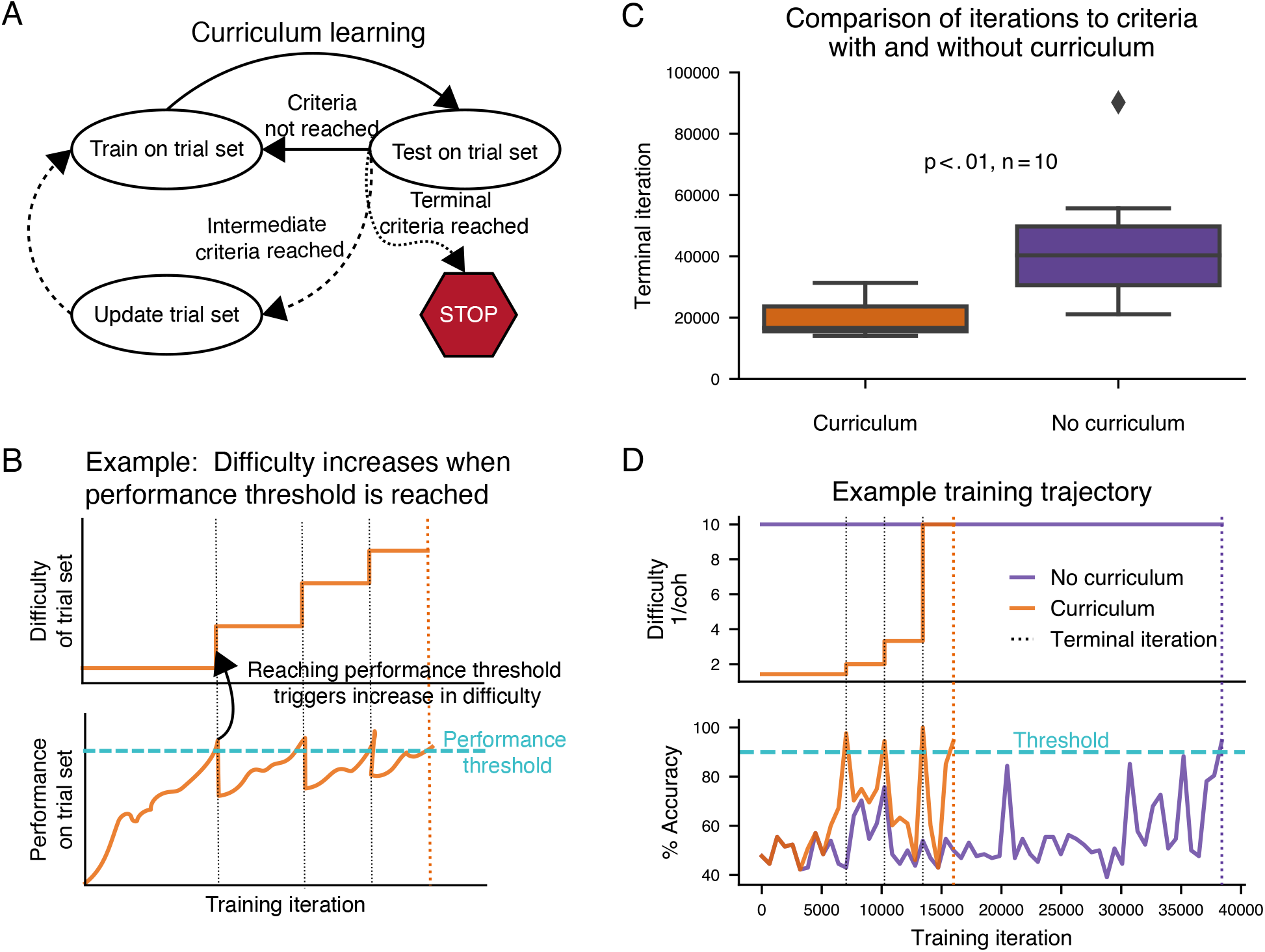
Curriculum learning. **(A)** Schematic of curriculum learning, or task shaping. The network is trained on selections from the trial set, then tested on selections from that trial set. Depending on the performance when testing on the trial set, the trial set can then be updated, e.g., to contain progressively more difficult trial conditions. **(B)** Example schematic of increasing difficulty of trial set (top) paired with performance over time (bottom). The task difficulty is progressively increased each time performance reaches the performance threshold. **(C)** Comparison of number of iterations needed to train a network to perform the perceptual discrimination task (from **Fig. 2**) with 90% accuracy at coherence level of 0.1. Ten networks were randomly initialized and each was trained both on a curriculum with decreasing coherence, and without a curricula with fixed coherence. Networks trained without curriculum learning were trained solely on stimulus with coherence = 0.1. Networks trained with curriculum learning were trained with a curriculum with coherence decreasing from 0.7 to 0.5 to 0.3 to 0.1 as performance improved (see **Extended Data, Figure 5-1**). When the network reached 90% accuracy on stimuli with coherence = 0.1, training was stopped. Networks trained with curriculum learning reached 90% accuracy significantly faster than networks without it (*p* < 0.01). **(D)** Trajectories of difficulty (defined here as inverse coherence), accuracy, and loss (mean squared error) across training iterations, for two identically initialized networks from **(C)**, one of which was trained with curriculum learning, and one of which was trained without curriculum learning.

### Comparison to other frameworks

PsychRNN compares favorably to alternative high-level frameworks available (**Table ??**). Most similar to PsychRNN is PyCog (Song et al., 2016), another Python package for training RNNs designed for neuroscientists. PsychRNN presents several key advantages over PyCog. First, PyCog’s backend is Theano, which is no longer under active support and development. Second, PyCog has no native implementation of curriculum learning. Third, task definitions in PyCog are not themselves modular, making experiments which are trivial to implement in PsychRNN more laborious and cumbersome for the user. Lastly, PyCog utilizes a built-in “vanilla” stochastic gradient descent algorithm, whereas PsychRNN allows users to select any optimizer available in the TensorFlow package.

Alternatively research groups may use a general purpose high-level wrapper of TensorFlow, such as Keras (Chollet and others, 2015), which is not specifically designed for neuroscientific research. Importantly, these frameworks do not come with any substantial ability to implement biological constraints. Users interested in testing the impact of such constraints would need to modify the native Keras Layer objects themselves, which is nontrivial. In addition, Keras does not provide a framework for modular task definition, which therefore requires the user to translate inputs and outputs into a form compatible with the model. PsychRNN, by close integration with the TensorFlow framework, manages to maintain much of the power and flexibility of traditional machine learning frameworks while also providing custom-built utilities specifically designed for addressing neuroscientific questions.

## Discussion

PsychRNN provides a robust and modular package for training RNNs on cognitive tasks, and is designed to be accessible to researchers with varying levels of deep learning experience. The separation into a Python- and NumPy-based Task object and a primarily TensorFlow-based Backend expands access to RNN model training without reducing flexibility and power for users who require more control over the precise setup of their networks. Further, the modularity of tasks and network elements enables easy investigation of how task and structure affect learned solution in RNNs. Lastly, the modular structure facilitates curriculum learning which makes optimization more efficient and more directly comparable to animal learning.

PsychRNN’s modular design enables straightforward implementation of curriculum learning to facilitate studies of how training trajectories shape network solutions and performance on cognitive tasks. Task shaping is a relatively understudied topic in systems neuroscience, despite its ubiquity in animal training. For instance, it is poorly understood whether differences in training trajectories result in different cognitive strategies or neural representations in a task (e.g., Latimer and Freedman, 2019). Standardization and automation in animal training may aid experimental investigation of task shaping effects (Murphy et al., 2020; Berger et al., 2018). PsychRNN provides an accessible framework to explore neuroscientific questions related to task shaping in RNN models.

Future extensions of the PsychRNN codebase can enable investigation into additional neuroscientific questions. Two potentially useful directions are the addition of units that exhibit firing rate adaptation through an internal dynamical variables associated with each unit (Masse et al., 2019) and the implementation of networks with short term associative plasticity (e.g., Miconi, 2017). An interesting area for extending task training capability is to add trial-by-trial dependencies. In the current version of PsychRNN, each task trial is trained independently from other trials in the same block. PsychRNN could potentially be extended to support dependencies across trials by having the loss function and trial specification depend on a series of trials.

The PsychRNN package provides an easy-to-use framework that can be applied and transferred between research groups to accelerate collaboration and enhance reproducibility. Where in the current environment research groups need to transfer their entire codebase in order to run an RNN model, in the PsychRNN framework they are able to transfer just a task or model file for researchers to investigate and build upon. The ability to test identically specified models across tasks in different groups, and identically specified tasks across models improves reliability of research. Furthermore, the many choices in defining and training RNNs can make precise replication of prior published research difficult. The specification of PsychRNN task files and parameter dictionaries can make reproduction of RNN studies more open and straightforward.

PsychRNN was designed to lower barriers to entry for researchers in neuroscience who are interested in RNN modeling. In service of this goal we have created a highly user-friendly, clear, and modular framework for task specification, while abstracting away much of the deep learning background necessary to train and run networks. This modularity also provides access to new research directions and a reproducible framework that will aid RNN modeling in neuroscientific research.

**Figure 6:**
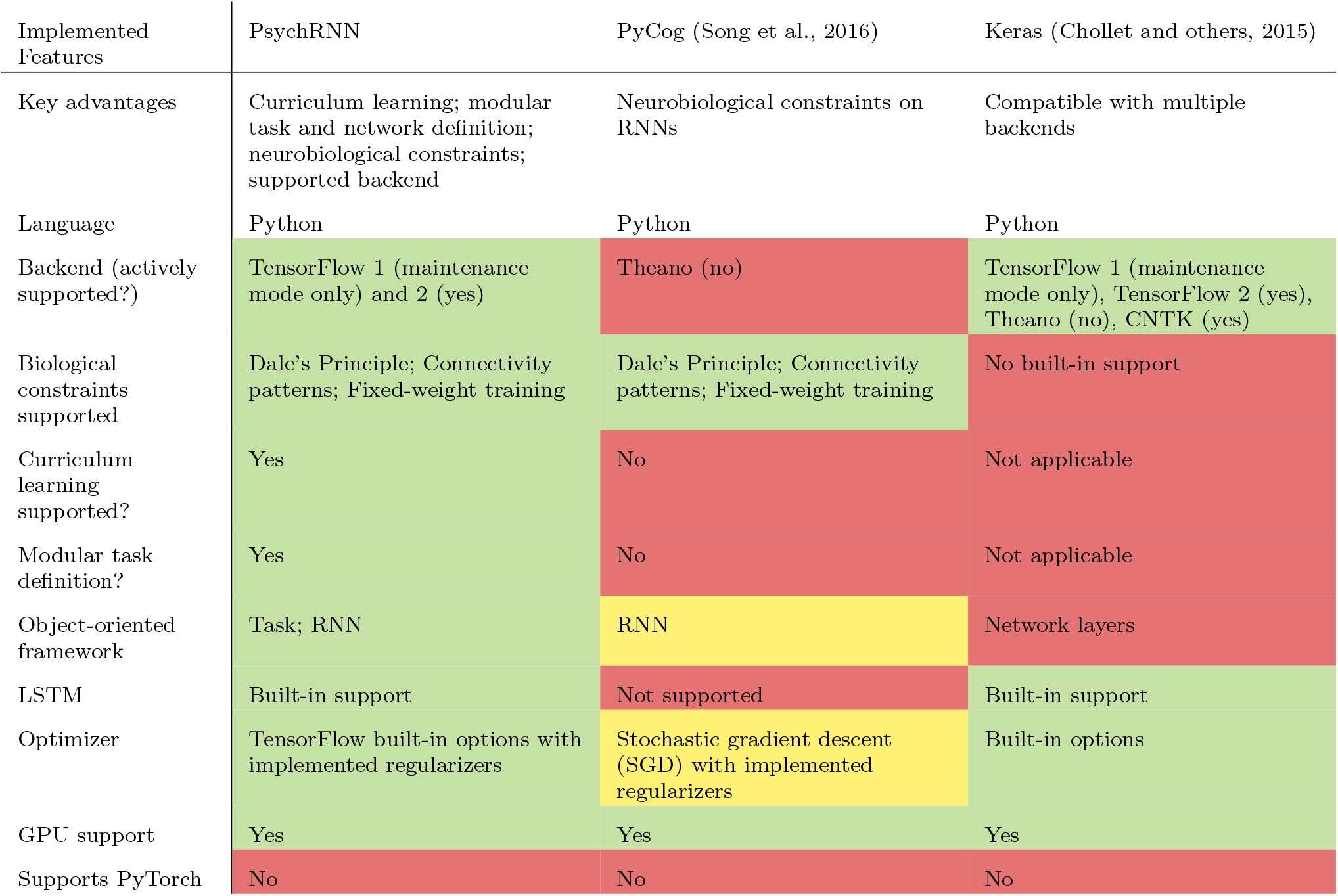
Comparison of PsychRNN to alternative RNN training packages. Red, yellow, and green indicate limited, moderate, and maximal flexibility or accessibility, respectively.

---

**Extended Data.** PsychRNN package and documentation.

**Extended Data, Figure 1-1:**
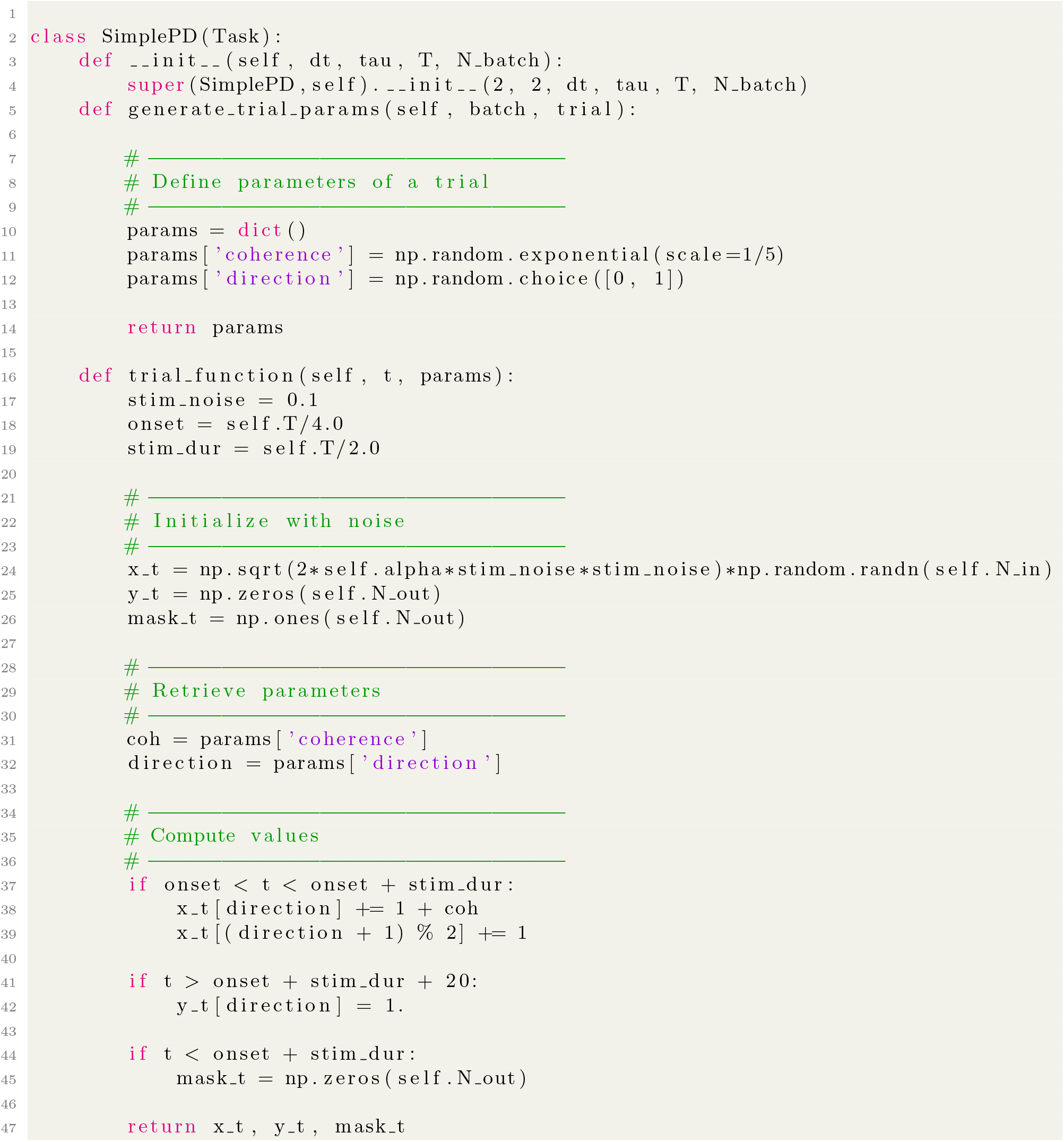
Example PsychRNN code showing task definition. This code sample defines a simple two-alternative perceptual decision-making task. The function `generate_trial_params` selects the coherence and direction on a trial by trial basis. `trial_function` sets the input, target output and output mask depending on the time in the trial and the passed parameters.

**Extended Data, Figure 3-1:**
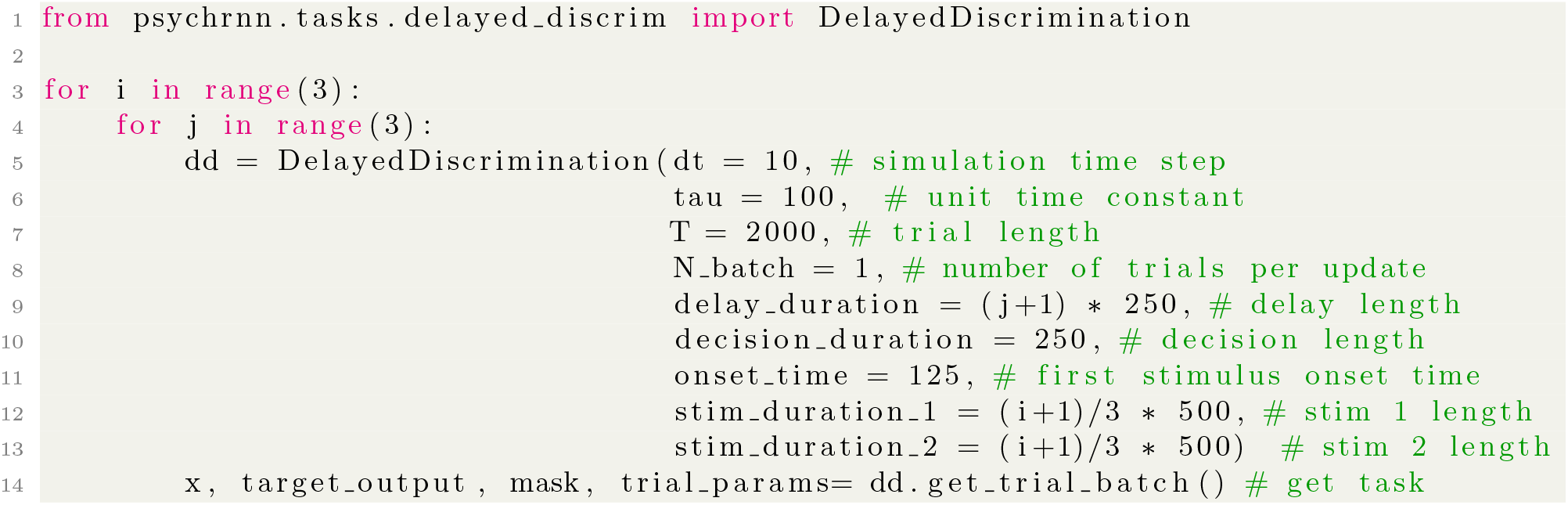
Example PsychRNN code showing modularity of task structure. This code sample produces all data shown in **Fig. 3B**. Although each plot in **Fig. 3B** has different task durations, iteration through all of them is simplified through the object-oriented modular task definitions enabled by PsychRNN. Resulting code is compact and readable compared to non-modular alternatives.

**Extended Data, Figure 3-2:**
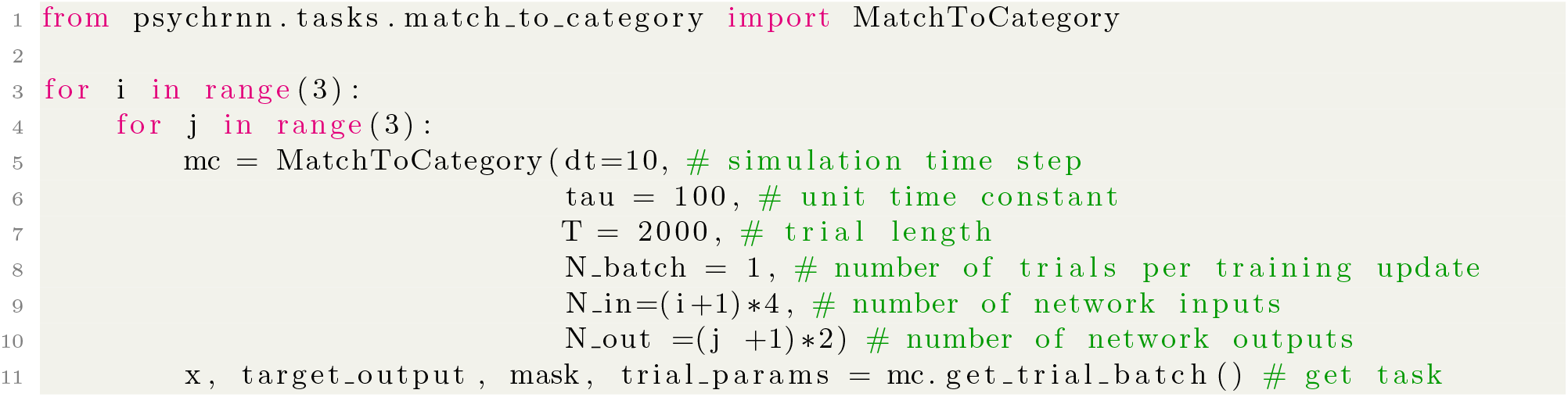
Example PsychRNN code showing modularity of task inputs and outputs. This code sample produces all data shown in Figure 3D. Although each plot in **Fig. 3D** has different numbers of inputs and outputs, iteration through all of them is simplified through the object-oriented modular task definitions enabled by PsychRNN. Resulting code is compact and readable compared to non-modular alternatives.

**Extended Data, Figure 5-1:**
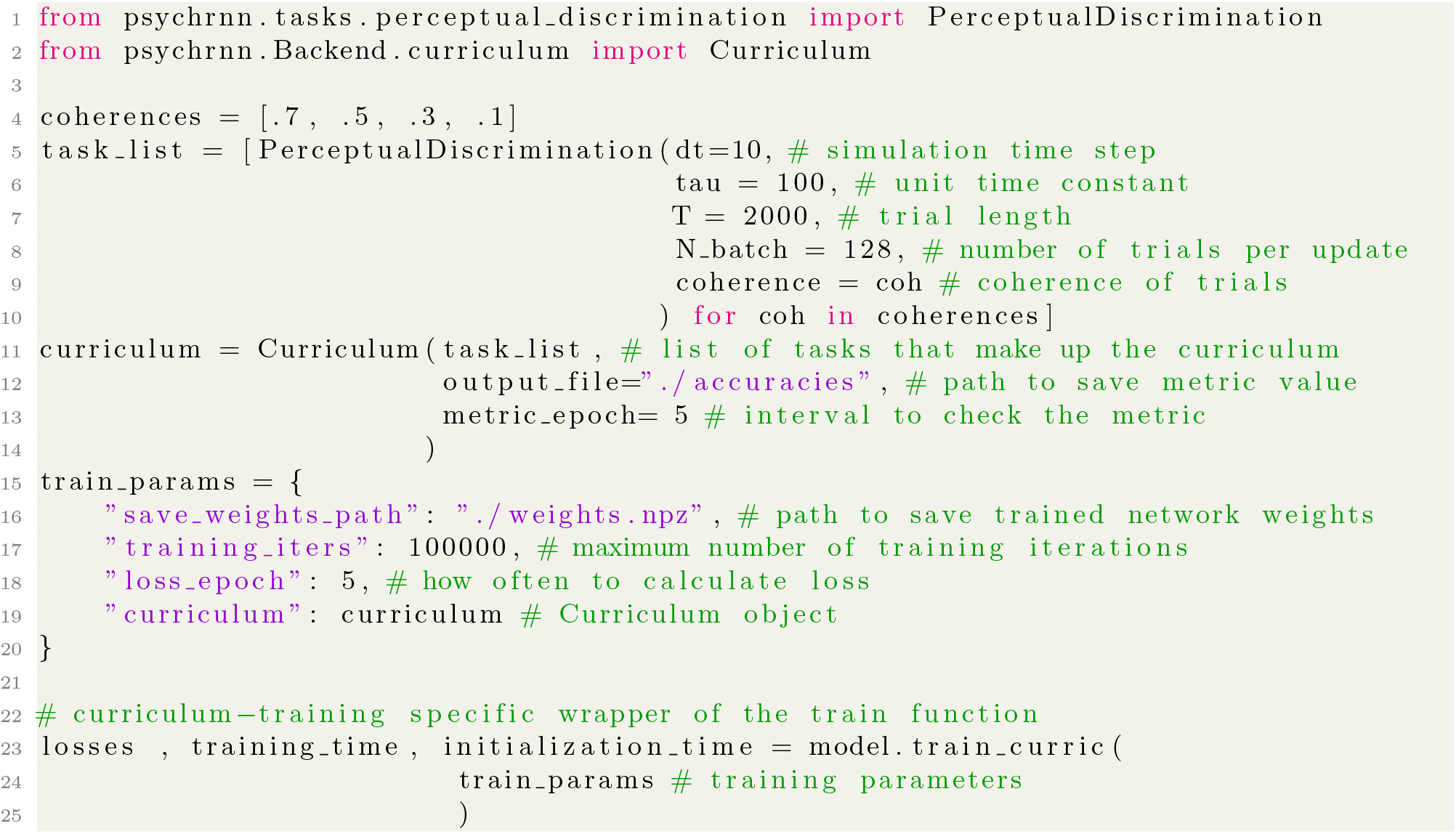
Example PsychRNN code showing curriculum learning. This code sample trains an RNN on a sequence of perceptual discrimination tasks with decreasing stimulus coherence. The network is first trained to perform the task with high coherence. Once the network reaches 90% accuracy on a given task (here, set of stimulus coherences), the network initiates training on the next task. This continues until the network has reached 90% accuracy on the final task—in this case, the lowest-coherence task. Curriculum learning is implemented by defining a list of tasks that form the curriculum (line 4-10), and passing that list in to the `Curriculum` class to form a `Curriculum` object (line 11-14). That `Curriculum` object is then included in the training parameters dictionary (line 15-20), and when the network is passed those training parameters for training, the network will be trained using the curriculum, or sequence of task parameters defined in lines 4-10.

